# Lactate-dependent metabolic rewiring is associated with CD163^+^ macrophage polarization in human pyelonephritis

**DOI:** 10.64898/2026.04.09.717269

**Authors:** Lars Borgards, Hannah Voss, Stephanie Tautges, Bente Siebels, Christoph Krisp, Devon Siemes, Philippa Spangenberg, Jenny Bottek, Sarah Scharfenberg, Jessica Schmitz, Lisa Schwarz, Christian Rehme, Boris Hadaschik, Jan H Braesen, Alexander Roesch, Hartmut Schlueter, Selina Jorch, Sibylle von Vietinghoff, Florian Wagenlehner, Olga Shevchuk, Daniel R. Engel

## Abstract

Chronic pyelonephritis is characterized by persistent bacterial infection of the kidney and a dysregulated immune response that promotes disease progression. Macrophages are central regulators of antibacterial immunity in the urinary tract. However, the mechanisms underlying their functional reprogramming in chronic infection remain poorly understood. Here, we identify lactate as a key metabolic determinant of macrophage polarization in human pyelonephritis. Proteomic analysis of human kidney tissue revealed extensive metabolic remodelling, including upregulation of enzymes involved in glycolysis. Consistent with this, lactate levels were significantly elevated in urine and plasma of pyelonephritis patients. Concomitantly, we observed a pronounced accumulation of CD163 macrophages in infected kidneys, representing a distinct macrophage subset with immunomodulatory function. Correlation-based network analysis revealed a strong association between CD163 and lactate dehydrogenase A, supporting a functional association between lactate metabolism and macrophage polarization. Mechanistically, exposure of murine bone marrow-derived macrophages to lactate induced intracellular protein lactylation and promoted polarization toward a CD163 phenotype, defining a metabolically imprinted macrophage state distinct from classical activation paradigms. Proteomic profiling demonstrated extensive remodelling of macrophage protein expression in response to lactate with significant alteration of mediators of phagocytosis, Toll-like receptor signalling, and interferon response. Together, these findings identify lactate as a potent metabolic driver of macrophage reprogramming and establish a foundation for investigating lactylation-dependent immune dysfunction in chronic pyelonephritis.

## Introduction

Macrophages are highly plastic immune cells that dynamically adapt their phenotype and function in response to local tissue cues, thereby shaping the outcome of bacterial infections^1,2^. In the urinary tract, macrophages orchestrate antibacterial defence, tissue repair, and resolution of inflammation through tightly regulated spatial and temporal programs^3-5^. However, persistent bacterial infections of the kidney, such as chronic pyelonephritis (PN), are frequently associated with dysregulated macrophage responses that favour immune suppression and disease persistence^6^. Among macrophage subsets, CD163^+^ macrophages are associated with anti-inflammatory, scavenging, and tissue-protective functions^7,8^. While these programs are beneficial during resolution phases, their sustained activation during ongoing bacterial infection may impair effective host defence. Indeed, CD163 expression is linked to immunoregulatory macrophage states across multiple inflammatory and infectious contexts^9^. However, the mechanisms that drive the accumulation and functional polarization of CD163^+^ macrophages in chronic kidney infection remain poorly understood.

Emerging evidence indicates that metabolic cues within inflamed tissues act as instructive signals for macrophage polarization^10^. Lactate, long considered a metabolic waste product, has recently been recognized as a signalling metabolite capable of reshaping immune cell function through receptor-mediated pathways, metabolic rewiring, and protein lactylation^11,12^. Histone lactylation has been shown to directly regulate gene expression programs associated with macrophage activation states^11^. However, whether lactate-driven macrophage reprogramming contributes to immune dysfunction and disease persistence in PN is currently unknown.

Lactate availability in macrophages is governed by the balance between glycolytic flux, mitochondrial pyruvate oxidation, and pyruvate-to-lactate conversion via lactate dehydrogenase A (LDHA)^13^. LDHA catalyses the conversion of pyruvate to lactate and thereby controls intracellular lactate pools that serve as substrates for protein- and histone lactylation. Importantly, modulation of LDHA activity alters lactate flux without directly impairing mitochondrial acetyl-CoA generation, which can be sustained through alternative metabolic pathways, such as amino acid and fatty acid metabolisms^13,14^. This metabolic architecture enables selective manipulation of lactate-dependent signalling while largely preserving global cellular viability and acetylation capacity. Our study identifies a lactate–dependent lactylation axis as a central mechanism linking metabolic tissue remodelling to macrophage-mediated polarisation in chronic PN.

## Results

To characterize metabolic alterations in human PN, we performed LC-MS/MS–based proteomic analysis of kidney tissue from PN patients and control donors. Principal component (PC) analysis of the total kidney proteome indicated PN-specific changes (Figure 1A). Pathway-enrichment analysis revealed coordinated upregulation of the innate immune response, whereas mitochondrial proteins involved in small molecule catabolism and mitochondrial ATP synthesis were downregulated, indicating substantial metabolic remodelling in infected kidneys (Figure 1B). Notably, enzymes associated with glycolytic flux were upregulated, suggesting a shift toward enhanced aerobic glycolysis (Figure 1C–D). Consistent with this metabolic phenotype, lactate concentrations were significantly elevated in both urine and plasma samples from PN patients compared to healthy controls (Figure 1E–F). These findings indicate that PN is characterized by a lactate-enriched systemic and local microenvironment.

**Figure 1.**
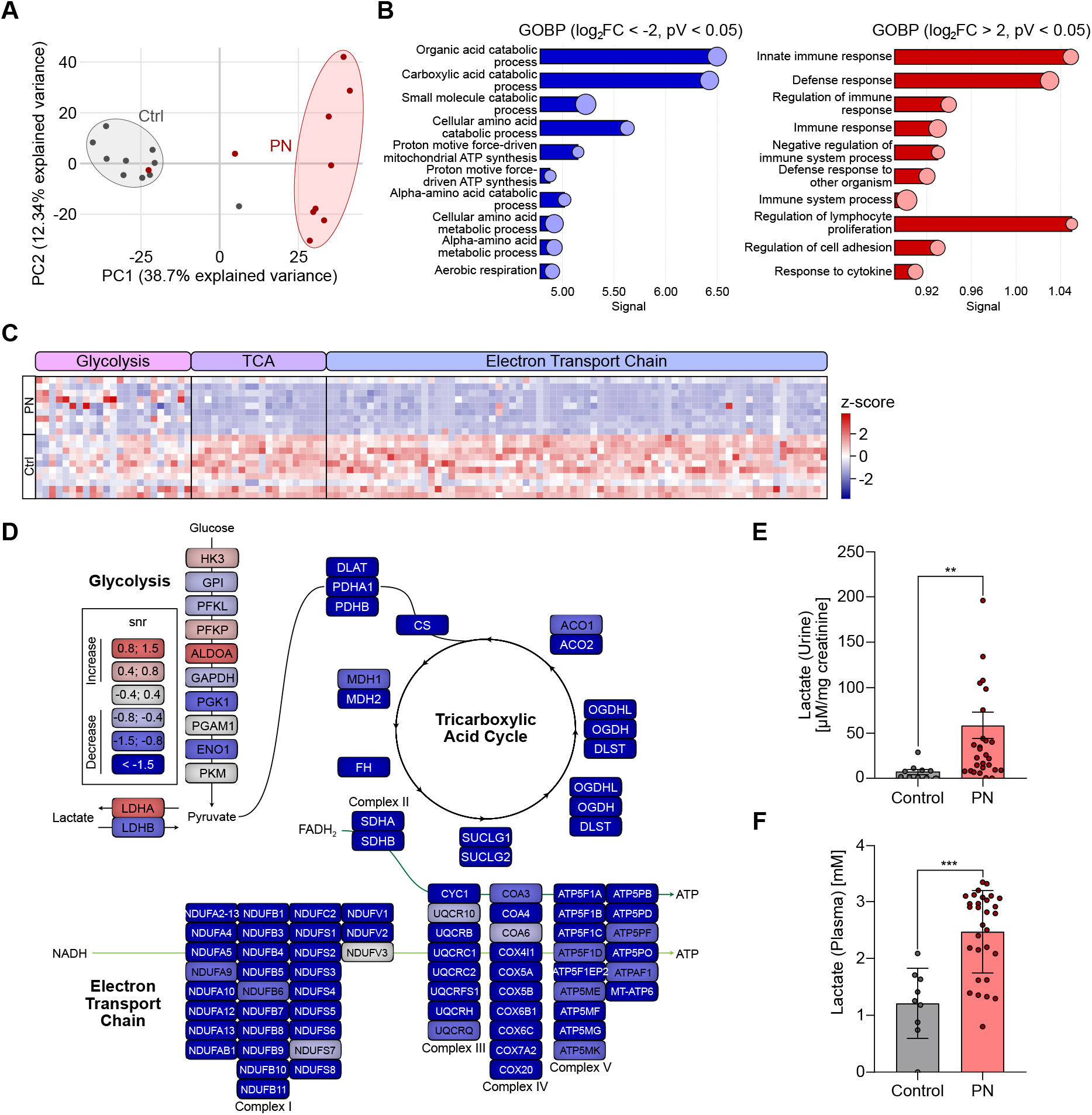
PN is associated with metabolic rewiring and increased lactate levels. **(A)** Renal tissue samples from PN (*n*=10) and control (*n*=10) patients were analysed by LC-MS/MS. Principal component (PC) analysis of the total kidney proteome indicated condition-specific changes. (**B**) Enrichment analysis of downregulated (blue) (log_2_(fold change) < -2, p-value < 0.05; left) and upregulated (red) (log_2_(fold change) > 2, p-value < 0.05; right) using the Gene Ontology Biological Process (GOBP) database. (**C**) Heatmap of z-score of protein groups annotated to glycolysis, the tricarboxylic acid (TCA) cycle and electron transport chain. **(D)** Schematic depiction of proteins involved in central carbon metabolism. Upregulation according to the signal-to-noise ratio (snr; PN vs. control) is indicated in red, downregulation in blue. **(E, F)** Lactate concentration was measured in urine (E) and plasma (F) samples from PN patients (*n*=30) and healthy donors (*n*=10). \*\**p*-value<0.01. \*\*\**p*-value<0.001.

To define molecular and spatial immune cell alterations in PN, we performed further proteomic and microscopy analysis. Among immune-associated proteins, CD163 emerged as one of the most strongly upregulated markers in PN tissue (Figure 2A). Immunohistochemical analysis confirmed a significant increase in CD163^+^ macrophages in infected kidneys compared to controls (Figure 2B–C). To further characterize these cells, we performed multiplex fluorescence microscopy, enabling simultaneous visualization of multiple immune cell populations, including markers of pro-inflammatory effector (CD11b, CXCR4, IFN-γ, iNOS), monocyte-derived (CD14), homeostatic/scavenging (CD11c, CD206, CD209, IRF4), and antigen-presenting cells (HLA-DR). CD68^+^ macrophages were abundant in the infected renal tissue with abundant frequency expressing CD163 (Figure 2D). Moreover, we observed a broad upregulation of functional markers, such as the pattern recognition receptor DC-SIGN (CD209), by CD163^+^ CD68^+^ macrophages (Figure 2E), supporting the concept that lactate environment in PN induce a distinct macrophage state with increased functional plasticity and transcriptional activity. Thus, these data identify CD163^+^ macrophages as a distinct and prominent immune cell population in human PN.

**Figure 2.**
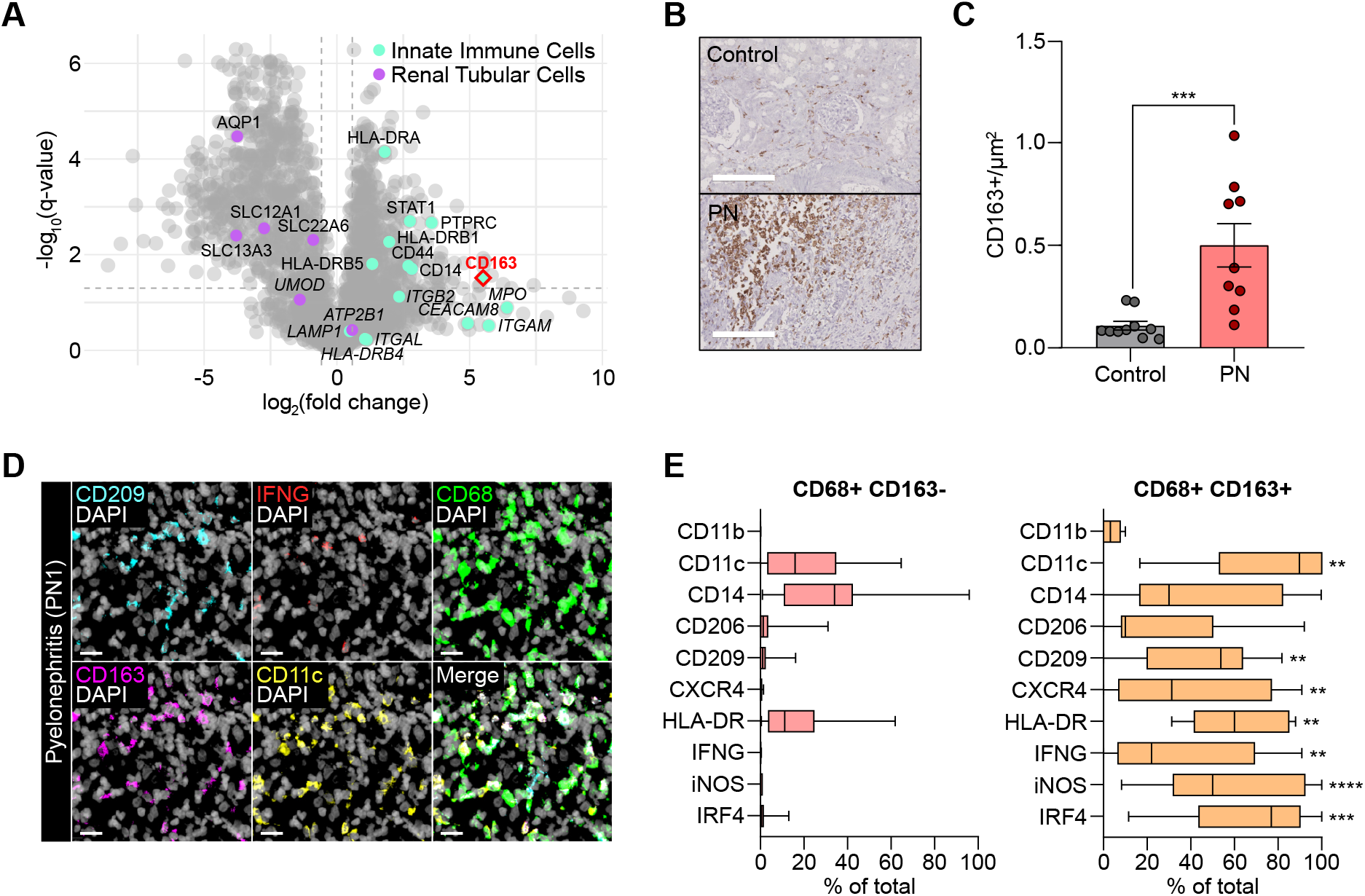
CD163^+^ macrophages represent a phenotypically distinct immune cell population enriched in PN. Volcano plot depicting the log_2_(fold change) and -log_10_*(q-*value) of proteins in PN vs. control as assessed by LC-MS/MS-based analysis of human kidney sections as indicated in Figure 1. Markers of innate immune and renal tubular cells are indicated in cyan and purple, respectively. **(B, C)** Representative images (B) and quantification (C) of the immunohistochemistry staining of CD163 in kidney tissue specimens. **(D, E)** Kidney tissue from PN patients (*n*=11) was analysed by multiplex fluorescence microscopy. (D) Representative images of CD68 (*green*), CD163 (*magenta*), and DAPI (*grey*). Scale bar represents 20 µm. E) Phenotypical analysis of multiplex microscopy data. Tukey box plots depict the percentage of CD68^+^ CD163^−^ and CD68^+^ CD163^+^ macrophages positive for polarization markers. \**p*-value<0.05, \*\**p*-value<0.01, \*\*\**p*-value<0.001, \*\*\*\**p*-value<0.0001

To investigate the relationship between macrophage polarization and metabolic remodelling, we performed correlation-based network analysis of the renal proteome. Proteins positively associated with CD163 abundance formed a coherent interaction network enriched for metabolic pathways (Figure 3A). Notably, lactate dehydrogenase A (LDHA), a key enzyme controlling lactate production, showed a strong positive correlation with CD163 expression across all analysed kidney samples, Pearson r >0.78 (Figure 3B). This association suggests that lactate metabolism is functionally linked to the emergence of CD163^+^ macrophages in the infected kidney.

**Figure 3.**
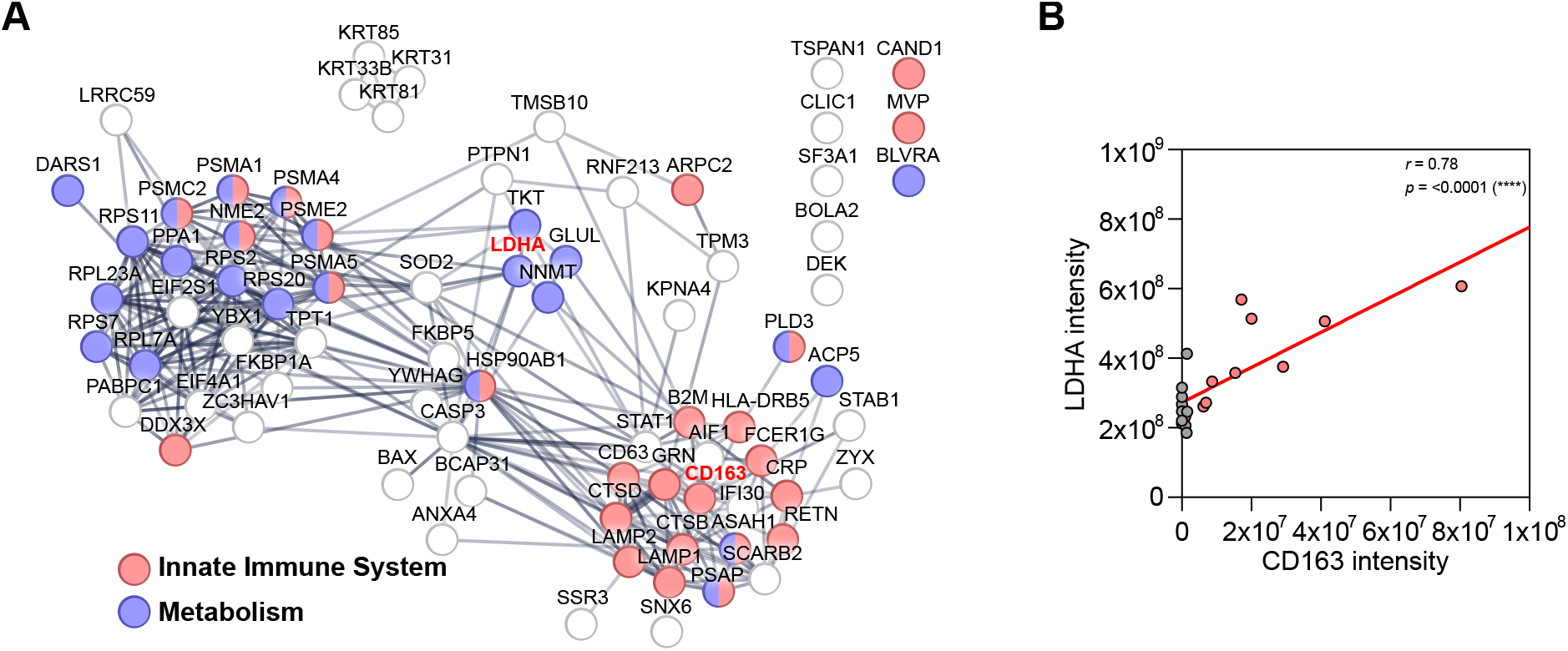
CD163 abundance correlates with lactate metabolism in human PN. **(A)** STRING network analysis of proteins detected by LC-MS/MS, which are positively correlated with the abundance of CD163 (Pearson r >0.6). Pearson correlation and linear regression analysis of the abundance of lactate dehydrogenase A (LDHA) vs. CD163. \*\*\**p*-value<0.001.

To determine whether lactate directly influences the macrophage phenotype, we performed *in vitro* experiments using murine bone marrow–derived macrophages (BMDMs). Cells were exposed to increasing concentrations of lactate to mimic the lactate-enriched environment in PN. Lactate treatment resulted in a significant increase in the proportion of CD163^+^ macrophages, accompanied by elevated intracellular protein lactylation as measured by flow cytometry (Figure 4A). Principal component analysis of ANOVA-regulated proteins quantified by LC-MS demonstrated clear separation of lactate-treated macrophages from classical pro-inflammatory (LPS/IFN-γ) and alternatively activated (IL-4/IL-13) cell states (Figure 4B). Proteomic analysis revealed extensive remodelling of these macrophages in response to lactate stimulation (Figure 4C). Differentially expressed proteins included regulators of immune cell identity, phagocytosis, TLR signalling, leukocyte migration, and cytokine production (Figure 4C). Upregulated proteins were enriched for factors of the response to interferon-α and -β, whereas regulators of the mitotic cell cycle and protein translation were decreased, indicative of terminally differentiated macrophages (Figure 4D). Together, these findings demonstrate that lactate induces a specific macrophage program, which is characterized by increased protein lactylation and polarization toward a CD163^+^ immunoregulatory phenotype.

**Figure 4.**
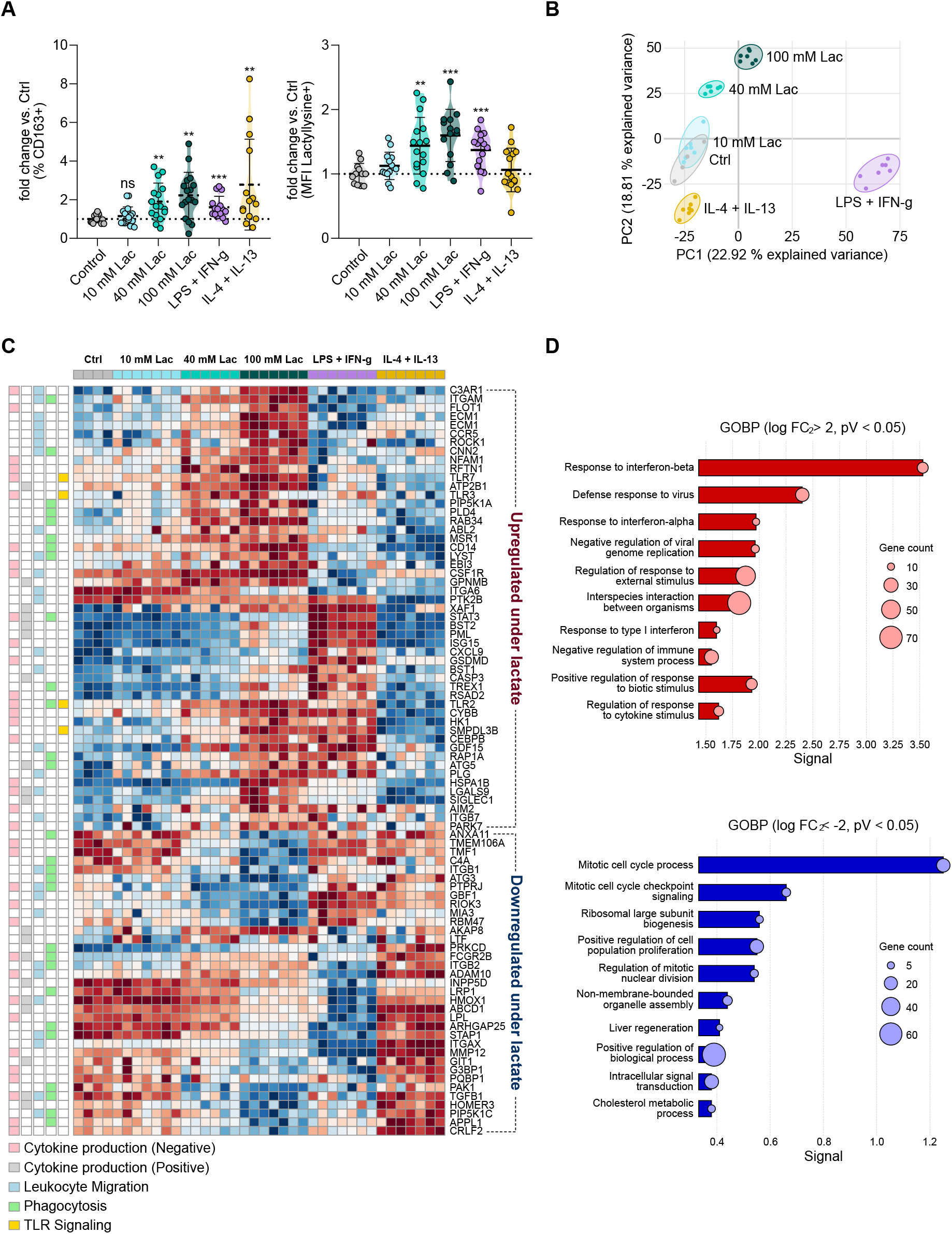
Lactate induces macrophage lactylation and drives CD163^+^ polarization. Murine BMDMs were treated with sodium-DL-lactate (Lac), LPS and IFN-g, or IL-4 and IL-13 for 48h. **(A)** Box plots depicting the fold change of the percentage of CD163^+^ cells (left) and the mean fluorescence intensity (MFI) of anti-lactyllysine (right), assessed by flow cytometry. The fold change was calculated for each replicate vs. the mean of the untreated control group. \**p*-value<0.05, \*\**p*-value<0.01, \*\*\**p*-value<0.001. **(B)** Principal component (PC) analysis of ANOVA-significantly (*p*-value<0.05) regulated proteins measured by LC-MS/MS proteomics. **(C)** Heatmap of log_2_-transformed abundance of proteins significantly regulated upon lactate treatment (*p*-value<0.05, fold change vs. control <-2 or >2) and annotated to negative (GO:0001818) or positive (GO:0001819) regulation of cytokine production, leukocyte migration (GO:0050900), phagocytosis (GO:0006909) or toll like receptor signalling pathway (GO:0002224). **(D)** Enrichment analysis based on the Gene Ontology Biological Process (GOBP) database of proteins significantly up-(top) and downregulated (bottom) upon lactate treatment.

## Discussion

In this study, we identify lactate as a central metabolic determinant of macrophage polarization in chronic PN and provide evidence that lactate-driven protein lactylation promotes the emergence of CD163^+^ macrophages with altered immunoregulatory function. By integrating human proteomics and spatial imaging, our findings establish a direct link between metabolic tissue remodelling and macrophage-mediated immune dysfunction in chronic kidney infection.

Our data demonstrate that PN is associated with profound metabolic rewiring characterized by enhanced glycolysis and elevated lactate levels. These findings are consistent with previous studies showing that inflamed and hypoxic tissues accumulate lactate due to increased glycolytic flux and altered mitochondrial function^14,15^. Importantly, lactate is increasingly recognized as an active signalling metabolite that regulates immune cell behaviour rather than merely reflecting metabolic stress^12^. A key finding of our study is the enrichment of CD163^+^ macrophages in PN tissue and their association with lactate metabolism. CD163^+^ macrophages have been widely described as anti-inflammatory and tissue-remodelling cells, often linked to resolution of inflammation and fibrosis^8^. However, in the context of persistent infection, such regulatory programs may become maladaptive and contribute to ineffective bacterial clearance. Our data extend this concept by identifying lactate as a potential upstream driver of CD163^+^ macrophage polarization in human PN.

Mechanistically, we show that lactate directly induces intracellular protein lactylation and drives macrophage reprogramming toward a CD163^+^ phenotype. These findings support the description of histone lactylation as a novel epigenetic modification linking metabolism to gene regulation in macrophages^11^. Subsequent studies have further demonstrated that lactylation can regulate inflammatory gene expression and immune cell differentiation^12,16^. Our results expand this framework by linking lactylation to impaired antibacterial effector functions in macrophages, including alterations in pathways related to phagocytosis, reactive oxygen species production, and cytokine responses.

Collectively, our findings position lactate not merely as a by-product of inflammation but as a central regulator of immune cell fate in chronic infection, which promote immune suppression and disease progression^17,18^. Thus, targeting lactate production, transport (e.g., monocarboxylate transporters), or lactylation pathways may represent a novel strategy to restore antimicrobial immunity in chronic PN.

## Materials Methods

### Kidney samples of PN patients

PN nephrectomy specimens (*n*=10; 20% men; mean age: 51.8 years; range: 22 – 76 years) and healthy regions from tumour resections (*n*=10; 30% men; mean age: 48.4 years; range: 7 – 82 years) were used from the archives of the Hannover Medical School (MHH). Ethic MHH: No. 10183_BO-K_2022. Additional fresh PN nephrectomy specimens (*n*=11; 36% men; mean age: 67.7, range: 46–76 years) were provided by the Department of Urology, University Hospital Essen (26-12914-BO).

### Urine and plasma samples of PN patients

Urine was collected from acute PN patients (*n*=30) and healthy control donors (*n*=10) by the Clinic for Urology, Pediatric Urology and Andrology of the Justus-Liebig-University Giessen. The ethical approval has been received by the independent ethics commission of the University Giessen, Germany (Ethical vote 280/20) on January 6^th^, 2021. The cases of acute PN were clinically diagnosed, following the FDA’s definitions used for studies in the indication “Developing drugs for treatment. Guidance for industry, June 2018” (https://www.fda.gov/media/71313/download). Patients admitted to the urological department at the University Hospital Giessen with at least two of the following signs or symptoms were included: chills, rigors, or warmth associated with fever; flank pain; nausea or vomiting; dysuria; urinary frequency; urinary urgency; costo-vertebral angle tenderness upon physical examination; or a urine specimen with evidence of pyuria. Risk factors for recurrent UTI were categorized by urologists according to the ORENUC classification system^19^. Urine samples were centrifuged at 5,000 × g for 10 min at 4 °C, and the supernatant was filtered through a 0.22 µm sterile filter (Merck Millipore, Burlington, USA).

### LC-MS/MS data acquisition and analysis of kidney tissue

FFPE kidney tissue samples were used for LC-MS/MS. Sample preparation and LC–MS/MS measurements were performed on a Orbitrap Fusion (Thermo Fisher Scientific) coupled to a nano-UPLC system (Dionex Ultimate 3000, Thermo Fisher Scientific) as previously described^20^. LC-MS/MS data were searched using the Sequest algorithm in Proteome Discoverer (v2.41.15, Thermo Fisher Scientific) against a reviewed human Swissprot database (April 2021; 20,365 entries). Carbamidomethylation of cysteine was set as a fixed modification, while methionine oxidation, N-terminal glutamine pyroglutamate formation, and protein N-terminal acetylation were included as variable modifications. Up to two missing tryptic cleavages were allowed, and peptides of 6 – 144 amino acids were considered. FDR < 0.01 was set for peptide and protein identification. Quantification was performed using the Minora Algorithm, followed by log_2_-transformation of protein abundancy and LOESS normalization^21^. For statistical analysis, 2,562 proteins detected in at least one condition with ≥2 biological replicates were included. For each protein, fold change, *p*-value, and signal-to-noise ratio were calculated, and *p*-values were adjusted for FDR using the Benjamini–Hochberg method (*q*-value). Proteins with |log_2_(FC)| > 2 and *q* < 0.05 were considered significantly regulated; 160 (6.2%) were upregulated and 507 (19.8%) downregulated. Cytoscape (Cytoscape_v3.8.0, ClueGO_v2.5.7; gene ontology databases from 08.05.2010; min GO level = 3; max GO level = 8; number of genes = 3; min percentage = 3.0; kappa score threshold = 0.4) was used for functional enrichment analysis.

### Measurement of lactate concentration in urine and plasma samples

Lactate concentration in human urine and plasma samples were determined using the Lactate-Glo− Assay (Promega) according to the manufacturer’s instructions. In short, samples were diluted 1:10 in PBS and 40 µL of diluted sample was transferred to a 96-well plate alongside titrated lactate standard. 40 µL of detection reagent were then added to each well and samples were incubated for 1h at RT. Luminescence was recorded on an iMark− plate reader (Bio-Rad). Urinary lactate concentrations were normalized to creatinine and differences in lactate concentration were tested for statistical significance by applying a Mann-Whitney U-test.

### Immunohistochemistry of CD163 and image analysis

Automated immunohistochemistry (Ventana Ultra; Ventana Medical Systems Inc.) was performed on FFPE human kidney tissue to stain CD163^+^ cells (Cell Marque, anti-human CD163 (MRQ-26) mouse monoclonal antibody). The number of CD163^+^ signals per mm^2^ was determined in FIJI and differences in the density of CD163^+^ cells between PN and control samples was tested for statistical significance by applying a Mann-Whitney U-test.

### Multiplex immunofluorescence microscopy by Akoya PhenoCycler− and image analysis

Fresh frozen human kidney tissue was sectioned at 10 µm thickness using a cryostat and mounted onto Superfrost− glass slides. Tissue was fixed with 4% paraformaldehyde for 10 min at RT, washed 1x with PBS and ddH_2_O, desiccated, and stored at 4°C until further use. To reduce tissue autofluorescence, slides were subjected to photobleaching prior to antibody staining. Here, sections were rinsed in ddH_2_O, and a bleaching solution (2% NaOH, 15% H_2_O_2_, 83% PBS) was applied. Slides were then exposed to broad-spectrum LED light for 45 min at RT. This step was repeated 1x with fresh bleaching solution. Following photobleaching, tissue sections were rinsed in ddH_2_O and washed 2x for 2 min each in Hydration Buffer, followed by incubation in Staining Buffer (Akoya Biosciences) for 20 minutes at RT. For multiplex immunofluorescence microscopy, all reagents were prepared according to standard laboratory protocols. A custom panel of DNA-barcoded antibodies compatible with the PhenoCycler Fusion platform (Akoya Biosciences) was used to characterize immune and metabolic features of the PN kidney. The panel comprised antibodies targeting CD68 (Akoya Biosciences, anti-human CD68 (AKYP0050)-BX015 monoclonal antibody), CD11b (Abcam, anti-human CD11b (EPR1344) rabbit monoclonal antibody), CD14 (Akoya Biosciences, anti-human CD14 (AKYP0079)-BX037 monoclonal antibody), CD163 (Akoya Biosciences, anti-human CD163 (AKYP0114)-BX069 monoclonal antibody), CD206 (R&D System, anti-human MMR/CD206 goat polyclonal antibody), CD209 (Abcam, anti-human DC-SIGN rabbit polyclonal antibody), CD11c (Akoya Biosciences, anti-human CD11c (AKYP0051)-BX024 monoclonal antibody), CD66b (BioLegend, anti-human CD66b (6/40c) mouse monoclonal antibody), HLA-DR (Akoya Biosciences, anti-human HLA-DR (AKYP0063)-BX033 monoclonal antibody), CD20 (Akoya Biosciences, anti-human CD20 (AKYP0049)-BX064 monoclonal antibody), IRF4 (RevMAb Biosciences, anti-human MUM1/IRF4 (RM352) rabbit monoclonal antibody), IFN-γ (Akoya Biosciences, anti-human IFNG (AKYP0093)-BX020 monoclonal antibody), CXCR4 (BioLegend, anti-human CD184 (12G5) mouse monoclonal antibody), MCT1 (Abcam, anti-human MCT1 rabbit polyclonal antibody), MCT4 (Abcam, anti-human SLC16A3/MCT4 rabbit polyclonal antibody), GLUT4 (Abcam, anti-human GLUT4 rabbit polyclonal antibody), iNOS (Akoya Biosciences, anti-human iNOS (AKYP0104)-BX023 monoclonal antibody), and lactyllysine (PTM Biolabs, anti-lactyl-L-lysine (9H1L6) rabbit monoclonal antibody). All antibodies were used at a dilution of 1:200. An empty barcode channel without primary antibody was included as a negative control to assess background signal. All antibodies were conjugated to unique DNA barcodes according to the manufacturer’s protocols and validated for use on fresh frozen tissue. A cocktail containing all DNA-barcoded antibodies and blocking reagents (N-, G-, J-, and S-blockers; Akoya Biosciences) was applied to each section and incubated overnight at 4 °C in a humidified chamber. After incubation, slides were washed thoroughly with Staining Buffer to remove unbound antibodies. After incubation with primary antibodies, samples were subjected to three consecutive fixation steps: 4% paraformaldehyde for 10 min, ice-cold methanol for 5min, and fixation reagent (1:50 in PBS) for 20 min, with intervening washes in PBS. Multiplexed imaging was performed using the PhenoCycler Fusion 2.0 system (Akoya Biosciences) according to the manufacturer’s instructions. Sequential imaging cycles were conducted by hybridizing fluorescently labelled oligonucleotides complementary to antibody barcodes, followed by imaging and signal removal. This iterative process enabled detection of all markers within a single tissue section. Images were acquired at high resolution using a 20× objective, with exposure settings optimized individually for each marker. Raw image data were processed using Akoya’s proprietary analysis software for cycle alignment, stitching, and background correction. Cell segmentation was performed based on nuclear staining (DAPI) using Cellpose-SAM^22^. Extraction of quantitative single-cell data and thresholding-based identification of cellular populations was performed in FIJI and “R” (v4.4.2, R Core Team (2021), respectively. Macrophages were identified as CD68^+^ CD66b^-^ CD20^-^ cells and separated into CD163+ and CD163-macrophages. Differences in the proportion of CD11b^+^, CD11c^+^, CD14^+^, CD206^+^, CD209^+^, CXCR4^+^, HLA-DR^+^, IFN-γ^+^, iNOS^+^, and IRF4^+^ subpopulations were tested for statistical significance by applying the Mann-Whitney U-test.

### Differentiation and stimulation of BMDMs

For the generation of M-CSF-enriched supernatant from L-929 mouse fibroblasts (DSMZ no.: ACC 2), cells were grown to confluency in T125 cell culture flasks in 100 mL of DMEM supplemented with 10% foetal calf serum (FCS), 1% penicillin/streptomycin (P/S), 1% L-glutamine, and 10 mM HEPES. Upon reaching confluency, cells were incubated for another 10 days. Afterwards, the supernatant was removed, sterile-filtered, and stored at -80°C until further use. Bone marrow cells were extracted from male adult mice (aged 8 – 10 weeks), passed through a cell strainer (40 µm pore size), centrifuged, and resuspended in RPMI 1640 supplemented with 10% FCS, 1% P/S, 1% L-glutamine, 10 mM HEPES, and 30% L-929-conditioned medium. 3 × 10^7^ bone marrow cells were incubated in 20 mL of differentiation medium in a Petri dish for 6 days. Upon differentiation, BMDMs were removed from the Petri dish by scraping, centrifuged, and resuspended in HPLM supplemented with 10% FCS, 1% P/S, 10 mM HEPES, and 50 µM β-Mercaptoethanol. 300,000 cells were seeded in 1 mL per well. BMDMs were treated with no stimulants (control), 10-, 40- or 100-mM sodium-DL-lactate, 15 ng/mL LPS and 50 ng/mL IFN-γ, and 20 ng/mL IL-4 and 20 ng/mL IL-13 for 48 h.

### Flow cytometric analysis of BMDMs

For flow cytometric analysis, BMDMs were washed 1x with 1 mL of PBS, detached by scraping, and transferred to 75 × 12 mm tubes. Fc receptors of suspended BMDMs were blocked with human immunoglobulin G (1:66 in PBS) for 10 min at 4°C. Cells were stained for F4/80 (Invitrogen, Alexa Fluor® 647 anti-mouse F4/80 (BM8) rat monoclonal antibody), CD11b (BioLegend, PE/Cyanine 7 anti-mouse/human CD11b (M1/70) rat monoclonal antibody), and CD163 (BioLegend, PE anti-mouse CD163 (S15049I) rat monoclonal antibody) for 20 min at 4°C in the dark. The LIVE/DEAD® Fixable Near-IR Dead Cell Stain Kit (Life Technologies) was used for staining of permeable, dead cells. Cells were then fixed with 4% paraformaldehyde and permeabilized using Perm/Wash Buffer (BD Biosciences) according to the manufacturer’s instructions. Intracellular lactylated proteins (PTM Biolabs, anti-L-lactyllysine rabbit polyclonal antibody) were stained for 20 min at 4°C in the dark. Incubation with secondary antibody (Abcam, Alexa Fluor® 488 anti-rabbit donkey polyclonal antibody) was performed for another 20 min at 4°C in the dark. Flow cytometric analysis was performed on a BD FACS Canto-II.

### LC-MS/MS sample preparation and data acquisition of BMDMs

For proteomic analysis, BMDMs were washed 3x with 1 mL of PBS each and lysed in 100 µL of lysis buffer (5% sodium-dodecyl-sulphate (SDS), 50 mM Tris, 150 mM NaCl, pH 7.8) supplemented with Complete Mini, EDTA-free protease inhibitor cocktail (Roche). Protein lysates were sonicated for 2 × 5 min and centrifuged at 5,000 g for 5 min. The supernatant was collected and stored at -80°C until further use. Protein concentration was determined using a bicinchoninic acid (BCA) assay (Pierce− BCA Protein Assay Kit, Thermo Fisher Scientific) according to the manufacturer’s instructions. For tryptic digestion, 2 µg of protein were reduced with 10 mM DDT at 56 °C for 30 min, followed by cysteine alkylation with 20 mM 2-iodoacetamide for 30 min at RT in the dark. Tryptic digestion was performed, using the single-pot, solid-phase-enhanced sample preparation (SP3) protocol, as previously described^23^. In brief, protein was bound to hydrophilic and hydrophobic beads (50:50 (w/w) using a bead to protein ratio of 10:1 for 18 minutes at RT. Samples were washed 2x with 100% acetonitrile (ACN) and 2x with 70% ethanol and reconstituted in 50 mM ammonium bicarbonate buffer. Tryptic digestion was performed at a trypsin-to-protein ratio of 1:100 for 16 hours at 37°C. Peptides were eluted with 20 µL 1% formic acid (FA) and 2% dimethyl sulfoxide in water. Peptides were vacuum-dried in a Concentrator plus (Eppendorf SE, Hamburg, Germany) and stored at −80 °C until further use. Directly prior to measurement, peptides were reconstituted in 0.1% FA. The peptide concentration was estimated using a NanoDrop Microvolume Spectrometer (Thermo Fisher Scientific, Waltham, USA).

### LC-MS/MS data acquisition and analysis of BMDMs

For proteomics, 300 ng of peptides in 0.1% FA were loaded onto an Evotip following the manufacturer’s instructions. Briefly, Evotips were activated with 0.1% FA in ACN, conditioned with 2-propanol, equilibrated with 0.1% FA, and then loaded for 1 min using centrifugal force at 800 g. Evotips were washed with 0.1% FA. Afterwards, 100 µL of 0.1% FA was added to each tip to prevent drying.

Peptide separation was performed on an Evosep One using the predefined 30SPD method, on an Aurora Elite CSI C18 UHPLC column (15 cm × 75 µm i.d., 1.7 µm, 120 Å; IonOpticks, Collingwood, Australia). Eluting peptides were injected into a TimsTOF^HT^ mass spectrometer, via a CaptiveSpray source at 1600V. The mass spectrometer was operated in data-independent acquisition (DIA) parallel accumulation-serial fragmentation (PASEF) mode. The accumulation and ramp time for the dual TIMS analyser were set to 100 ms at a ramp rate of 9.42 Hz. Scan ranges were set to m/z 100–1,700 for the mass range and 1*K0-1 0.60–1.60 V s −1 cm −2 for the mobility range at both MS levels. For DIA-PASEF, a cycle time of 1.8 seconds was estimated. Within each cycle, a precursor mass range of 400 to 1201 Da and a mobility range of 0.6 to 1.6 1 K0-1 was fragmented. At MS1 level, one ramp was executed per cycle. For MS/MS measurements 32 MS/MS windows with a mass width of 26.0 Da each and a mass overlap of 1.0 were distributed across the 16 MS/MS ramps. LC-MS raw data were processed in DIA-NN (v1.9.2, Aptila Biotech^24^), applying the following settings: Enzyme/Cleavage rules: Trypsin/P; Fixed modification: Carbamidomethyl (C); Variable modifications: Acetyl (Protein N-term) and Oxidation (M)). Raw data were searched against a reviewed murine UniProt/Swissprot FASTA database (Version 25 February 2025, containing 17221 target sequences). Peptides and proteins were considered identified with an FDR < 0.01 at peptide and protein level, respectively. Quantification was performed at the MS2 level, with RT-dependent normalization enabled. Protein abundances were exported and log_2_-transformation as well as imputation of missing values from normal distribution were performed in Perseus (v2.1.1.0). Further data analysis was performed in the R software environment (v 4.4.2, R Core Team (2021)). Here, differentially abundant proteins between pairs of predefined phenotypes were determined through Student’s t-testing. Proteins, identified with a *p*-value < 0.05 and at least 2-log_2_(fold change) difference between groups, were considered significantly regulated and subjected to further enrichment analysis using the STRING database (v12.0).

## Author contributions

L.B., B.Si., S.T., J.B., S.S., L.S., J.S. performed experiments

L.B., P.S., D.S., and H.V. analysed data.

J.S., H.H.B., C.C, H.S., S.V.V., C.R., B.H., A.R., F.W. provided resources, reagents, and expertise.

L.B., O.S. and D.R.E. wrote the original draft.

D.R.E. and O.S. supervised the work, acquired funding, provided project administration, and conceived and designed the study.

All authors contributed to review and editing of the manuscript. No conflict of interest exists.

## Acknowledgements

We acknowledge the following support by the University Duisburg Essen: Open Access Publication Fund, the Central Animal Facility, the Imaging Center Essen, the MOSAIC Facility, the department of Experimental and Translational Research, the department of Neutrophil Biology and Translational Oncology and the Immunoproteomics group supported by INST 20876/486-1. We received funding from the German Research Foundation: FOR5427 SP1-SH542-3/1, ERA-NET NEURON (01EW2503) (OS); FOR5427 (466687329) to SP1 (FW), SP4 (DRE), SP6 (SJ), SP7 (SVV); EN984/16-1, 18-1 and 19-1 (DRE); TR332 (449437943) A3 and Z1 (DRE), INST 20876/486-1 (DRE), 516868494 (HS), 518551069 (HS), 247354600 (HS), 247377969 (HS), 426788273 (HS) and the Mildred Scheel Cancer Career Center Hamburg (JH). MALDI-MSI measurements were supported by Bruker Daltonics.

## Notes

### Competing Interest Statement

The authors have declared no competing interest.

